# Synthesis of modified nucleotide polymers by the poly(U) polymerase Cid1: Application to direct RNA sequencing on nanopores

**DOI:** 10.1101/2021.07.06.451372

**Authors:** Jenny Vo, Logan Mulroney, Jen Quick-Cleveland, Miten Jain, Mark Akeson, Manuel Ares

## Abstract

Understanding transcriptomes requires documenting the structures, modifications, and abundances of RNAs as well as their proximity to other molecules. The methods that make this possible depend critically on enzymes (including mutant derivatives) that act on nucleic acids for capturing and sequencing RNA. We tested two 3’ nucleotidyl transferases, *Saccharomyces cerevisiae* poly(A) polymerase and *Schizosaccharomyces pombe* Cid1, for the ability to add base and sugar modified rNTPs to free RNA 3’ ends, eventually focusing on Cid1. Although unable to polymerize ΨTP or 1meΨTP, Cid1 can use 5meUTP and 4thioUTP. Surprisingly, Cid1 can use inosine triphosphate to add poly(I) to the 3’ ends of a wide variety of RNA molecules. Most poly(A) mRNAs efficiently acquire a uniform tract of about 50 inosine residues from Cid1, whereas non-poly(A) RNAs acquire longer, more heterogeneous tails. Here we test these activities for use in direct RNA sequencing on nanopores, and find that Cid1-mediated poly(I)-tailing permits detection and quantification of both mRNAs and non-poly(A) RNAs simultaneously, as well as enabling the analysis of nascent RNAs associated with RNA polymerase II. Poly(I) produces a different current trace than poly(A), enabling recognition of native RNA 3’ end sequence lost by in vitro poly(A) addition. Addition of poly(I) by Cid1 offers a broadly useful alternative to poly(A) capture for direct RNA sequencing on nanopores.

## INTRODUCTION

RNA plays dual roles during gene expression, both carrying information and interpreting it along the way. RNAs containing protein coding information from genes are acted upon by non-coding RNAs during mRNA processing, translation, and decay. Correct reformulation and use of primary gene transcripts in context critically depends on intricate enzymes with non-coding RNA subunits: the spliceosome, ribosome, and miRNA complexes among others. The composition of cellular RNA populations, both coding and non-coding, reflect cell state. Tracking these states over time can reveal what a cell has done, what it is doing at the moment, and what it might soon do. Rapidly evolving methods for high throughput RNA sequencing have diversified and multiplied to the extent that our ability to detect and count RNAs by their structure, modification status, location, and associations with other cell components is limited only by the RNA populations we can capture (Pachter (2013). In turn the development of these technologies has relied on foundational research into the capabilities of proteins that manage nucleic acid transactions, and the application of these enzymes in innovative ways.

One such useful enzymatic activity is template-independent nucleotide addition to the 3’ ends of RNA and DNA. The discovery and use of polynucleotide phosphorylase to create RNA polymers can be traced to the origins of molecular biology (Grunberg-Manago et al. 1956; Grunberg-Manago 1989). Terminal deoxynucleotidyl transferase played a key role in early cloning strategies (Jackson et al. 1972), and more recently applications of poly(A) polymerases for labeling or tailing of RNA have been developed (Winter and Brownlee 1978; Lingner and Keller 1993; Martin and Keller 1998). As RNA 3’ nucleotidyl transferases have been studied more broadly, structurally related enzymes that add uridine as well as other nucleotides have been characterized (Liudkovska and Dziembowski 2021; Preston et al. 2019; Shukla et al. 2020), suggesting that a variety of homopolymeric or mixed polymer 3’ tails other than poly(A) could be created for different purposes, including bar-coding. Although many studies have tested for 3’ nucleotidyl transferase activity using the four standard ribonucleotides, few studies have addressed the extent to which modified or unnatural bases can be used as substrates. In general, understanding the ability of 3’ nucleotidyl transferases to create non-native homogeneous or mixed polymer 3’ tails could lead to technological applications that might enable further discovery.

A technology arena that awaits improvements is direct RNA sequencing using nanopores, implemented commercially by Oxford Nanopore Technologies, Inc. (Workman et al. 2019; Garalde et al. 2018). Advantages of direct RNA sequencing by nanopores include the ability to acquire information from full-length single molecules that is lost during the fragmentation necessary for short-read sequencing. Perhaps more intriguing is the potential to read modified nucleotides (Smith et al. 2019; Leger et al. 2019; Begik et al. 2021), whose presence is usually erased by reverse transcription in DNA-based RNA sequencing methods. Despite these advantages, direct RNA sequencing on nanopores suffers from low throughput and limited ability to pool samples within the same sequencing run. Although custom adapters can allow individual RNAs to be targeted (e.g. Smith et al. 2019), the commercial method targets poly(A)+ mRNAs, leaving most non-polyadenlyated noncoding RNAs (ncRNAs) inaccessible (Workman et al. 2019; Garalde et al. 2018).

Our interest in accessing sequence and modification status for more of the transcriptome led us to propose that addition of unnatural or rare nucleotides to the 3’ ends of RNA could both allow broad capture of more diverse populations of RNA without loss of native 3’ end information. Furthermore, distinct homopolymers or mixed polymers offer the potential for barcoding and pooled sequencing of multiple samples on nanopores. To this end we explored the ability of available 3’ nucleotidyl transferases to incorporate nucleotides other than the standard four that make up most transcripts in the cell. Here we document several novel in vitro activities of the Cid1 poly(U) polymerase (Rissland et al. 2007), including the ability to add long inosine tails to the 3’ ends of a wide variety of RNAs. For unknown reasons Cid1 addition of inosine to polyadenylated mRNAs stalls after addition of 50 residues. Addition of modified U residues is also unusual: 5methylUTP and 4thioUTP are readily incorporated, but ΨTP and 1meΨTP are not. Finally we demonstrate the application of the novel inosine activity in direct RNA sequencing on nanopores, enabling measurement of both mature and nascent mRNA populations as well as noncoding RNAs such as telomerase, spliceosomal snRNAs, snoRNAs, RNaseP RNA, and others.

## RESULTS

### Cid1 poly(U) polymerase can efficiently add inosine to the 3’ ends of RNA

To find new ways to add modified nucleotides to the 3’ ends of RNA, we tested commercial preparations of two well-studied enzymes, the *S. pombe* poly(U) polymerase Cid1 (Rissland et al. 2007), and *S. cerevisiae* poly(A) polymerase (PAP) (Martin and Keller 1998). We first confirmed that the commercial preparations had the expected nucleotide adding specificities by incubating them with various rNTPs and a 24-nucleotide oligomer of adenosine (A24) under the reported conditions (Fig 1A and 1B). Briefly, Cid1 from New England Biolabs efficiently adds U and A, but only poorly adds C or G (Fig 1A), in agreement with previous work (Rissland et al. 2007). PAP from ThermoFisher efficiently adds A, and to a much lesser extent G, but only poorly adds U or C, as observed previously (Martin and Keller 1998). We next tested whether the enzymes could use inosine triphosphate (ITP, a purine similar to G) to make poly(I) tails. Cid1 added long stretches of inosine to A24 (Fig 1A, lane 6) whereas PAP only added a few residues (Fig 1B, lane 6). Addition of inosine by Cid1 was unexpected since the enzyme does not use GTP very well (Rissland et al. 2007, Fig 1A) and inosine differs only by the lack of the 2-amino group present on guanosine.

**Figure 1.**
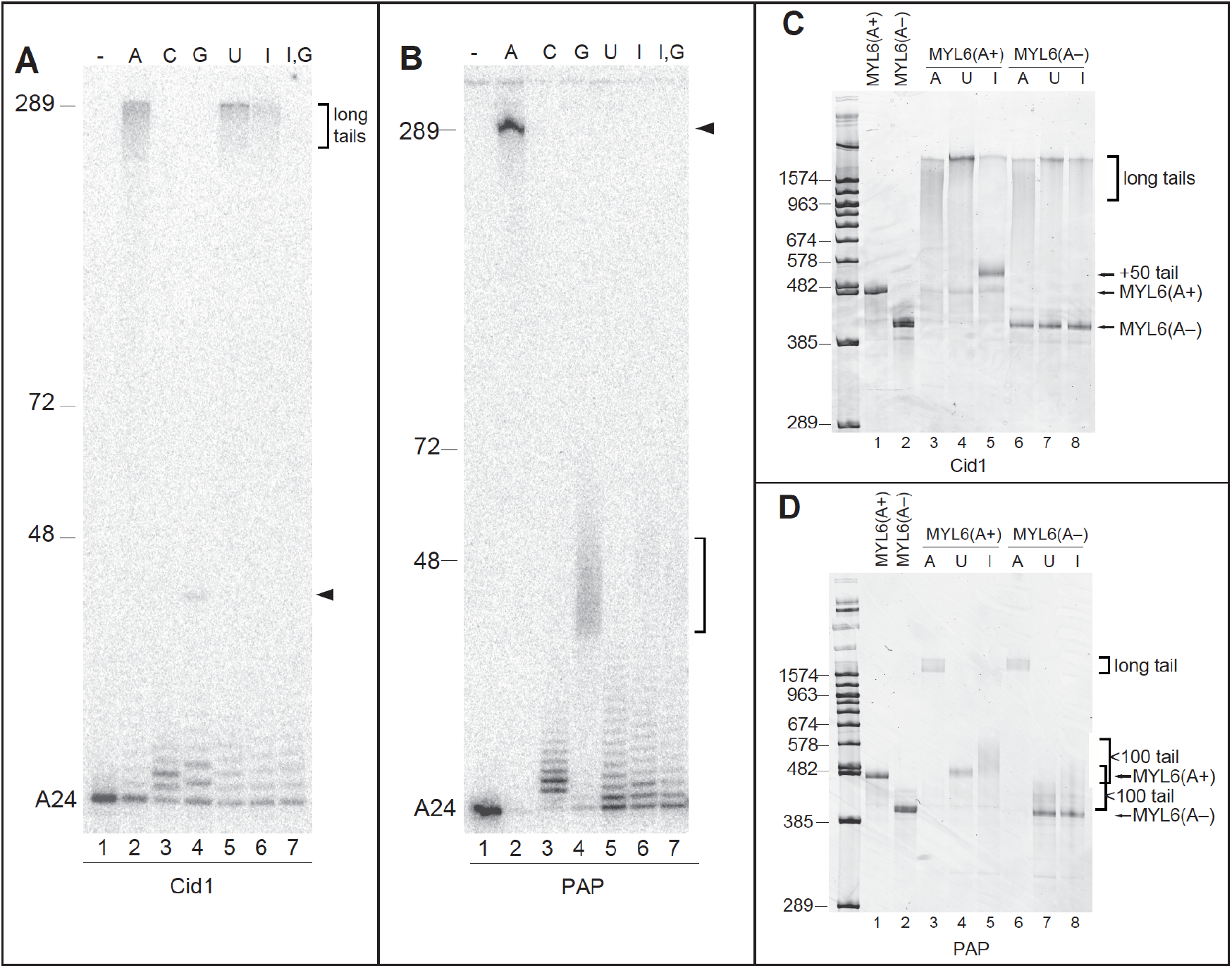
Nucleotide-adding activity of *S. pombe* Cid1 and *S. cerevisiae* Poly(A) polymerase (PAP) on various model RNA substrates. RNA was incubated with enzyme and rNTPs as indicated and run on denaturing polyacrylamide gels, stained with SYBR Gold and imaged on a Typhoon scanner. Markers are DNA with the indicated chain lengths for the equivalently migrating RNA, using the ~1.04X greater mass to charge ratio of RNA per residue. For example, a 500 nt DNA marks the approximate migration of a 482 residue RNA. **A.** Cid1 adds long tails of A, U, or I, but much shorter tails of C or G to A24. **B.** PAP adds long tails of A, a 35-50 nt stretch of G, but only short stretches of C, U, or I to A24. **C.** Cid1 efficiently adds long stretches of A or U, but mostly just 50 residues of I to a model poly(A)+ mRNA, whereas it inefficiently adds A, U, or I to a non-polyadenylated RNA. **D.** PAP efficiently adds A to either polyadenylated or non-polyadenylated RNA (lanes 3 and 6) and adds a short stretch of I to poly(A)+ RNA, but only inefficiently adds U or I to non-polyadenylated RNA.

To test substrates more representative of natural mRNAs, we prepared model mRNAs based on human MYL6 with or without a 40 nucleotide poly(A) tail, to represent polyadenylated (MYL6(A+)) and non-polyadenylated (MYL6(A-)) RNAs. As with the A24 substrate, Cid1 used UTP or ATP to generate long poly(U) or poly(A) tails on both model RNAs (Fig 1C, see also Rissland et al. 2007; Yates et al. 2012; Lunde et al. 2012; Munoz-Tello et al. 2012). Unexpectedly, Cid1 uniformly added about 50 inosine residues to a majority of MYL6(A+) molecules, but also produced a few molecules with long (>200 nt) tails (Fig 1C, lane 6). In contrast, Cid1 inefficiently added long tails on a fraction of MYL6(A-) molecules (Fig 1C, lane 8). The low efficiency of Cid1 addition of U, A, or I to this molecule (compare lane 8 with lanes 6 and 7) suggests poor access to the 3’ end of the substrate RNA rather than a preference for a particular rNTP substrate.

In comparison, PAP efficiently added poly(A) to both MYL6(A+) and MYL6(A-) RNAs, but only poorly added U to either (Fig 1D, compare lanes 3 and 4 to lanes 6 and 7, see also Martin and Keller 1998). The presence of a poly(A) tail on MYL6(A+) allowed PAP to add short heterogenous stretches of inosine to most of the molecules (Fig1D, lane 5), but MYL6(A-) is much less efficiently used (Fig 1D, lane 8). When we tested Cid1 and PAP using a shorter (200 nt) model mRNA with 44 A residues (Gluc200(A+)) or without a poly(A) tail (GLuc200(A-)), we saw comparable activities except that, similar to its activity on the A24 substrate (Fig 1A, lane 6), Cid1 did not stop at +50 on Gluc200(A+) (Suppl Fig S1A, lane 5). This suggests that a starting substrate RNA of a certain length, or with a certain length of poly(A), may be necessary to create the +50 product. We conclude that the presence of a pre-existing poly(A) tail significantly promotes the use of an RNA as a substrate for both enzymes, except that RNAs below a certain size or with other as yet unidentified structural features (such as an inaccessible 3’-OH) may be poor substrates.

### Validation of inosine addition by Cid1

To precisely measure the lengths of the poly(I) extension of RNA catalyzed by Cid1, we labeled various I-tailed and untailed control RNAs at their 3’ ends with ^32^P-pCp and T4 RNA ligase. We then digested the labeled RNA with RNAse A, which cuts only after pyrimidines, leaving homopurine polymers like poly(A) or poly(I) intact and still carrying the radioactive phosphate (Fig 2A). For example, MYL6(A+) contains a U followed by two Gs at its 3’ end to which the template encoded 40 nucleotide poly(A) tail is added during T7 transcription. Thus the MYL6(A+) substrate labeled in this way is predicted to have a 43 nt RNAse A-resistant poly(A) containing oligomer GGA_40_*pCp (* indicates the radioactive phosphate, designated as GGA_40_ in Fig 2B). In the I-tailed MYL6(A+) digest, the 3’ end product is ~93 nucleotides, confirming that Cid1 adds ~50 inosine residues onto the pre-existing poly(A) tail. The non-polyadenylated RNAs MYL6(A-) and yeast 5.8S rRNA acquire a heterogeneous I-tail that can be much longer than 50 residues (Fig 1C). Accordingly, RNase A digestion of each of these ^32^P-pCp labeled Cid1 products generates a ladder of RNAse A-resistant products that extends far up the gel (Figs 2C and D). We conclude that Cid1 generally adds a uniform ~50 nt I-tail to poly(A)+ RNAs, whereas it adds from a few to >1000 inosines to non-polyadenylated RNAs.

**Figure 2.**
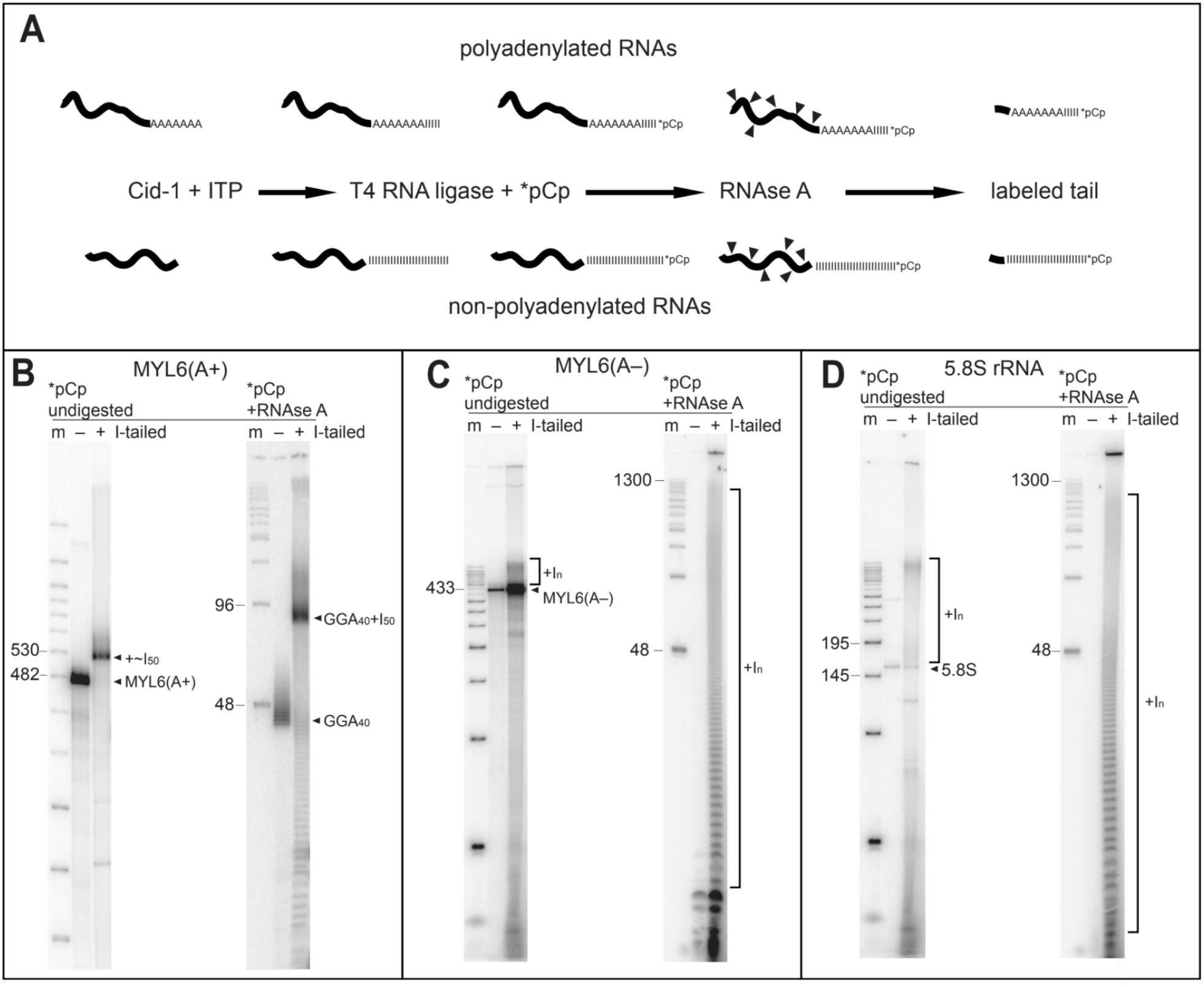
Biochemical determination of inosine tail lengths. **A.** Scheme for analyzing inosine tails based on RNAse A resistance of purine homopolymers. Top, poly(A)+ mRNAs, bottom, non-polyadenylated RNAs. From left to right: tailing by Cid1, 3’ end labeling by T4 RNA ligase and ^32^P-pCp, RNAse A digestion. **B.** Cid1 adds ~50 inosine residues onto the poly(A) tail of a model poly(A)+ mRNA. Left, labeled products without (-) and with (+) I-tailing, and without RNAse A digestion, run on a 6% denaturing polyacrylamide gel. Right, labeled products without (-) and with (+) I-tailing after RNAse A digestion, run on a 12% denaturing polyacrylamide gel. Markers are indicated as for Fig 1. **C.** Cid1 adds a long heterogeneous tract of inosine residues onto the 3’ end of a model non-polyadenylated mRNA. Lanes are as for (B) above. **D.** Cid1 adds a long heterogeneous tract of inosine residues onto the 3’ end of the non-polyadenylated 5.8S rRNA. Lanes are as for (B) above.

To confirm that addition of the ~50 I-tail is an intrinsic feature of Cid1 and not generated by an unknown step in the commercial preparation of the enzyme at New England Biolabs, we cloned, expressed, and purified a recombinant Cid1 with a truncated N-terminus (vCID1) in *E. coli* (see Supplemental Methods). We found that the addition of ~50 inosines to MYL6(A+) is a property of both Cid1 preparations and is not dependent on unknown commercial purification or treatment steps (Suppl Fig S2A).

### Testing the ability of Cid1 to use modified uridine triphosphates

Given its native uridylation activity, we also tested the ability of Cid1 to incorporate modified U residues (Fig 3), both alone (at 1 mM) and in combination with unmodified UTP (0.5 mM each). We find that Cid1 is unable to add long tracts of 2’-O-methyl-UTP to RNA (Fig 3, lane 3), and is blocked in the addition of UTP when 2’-O-methyl-UTP is present in the same reaction (lane 4), suggesting that incorporation of 2’-O-methyl-UTP is chain terminating for the Cid1 reaction. Cid1 is also unable to add ΨTP efficiently, however ΨTP does not greatly inhibit incorporation of UTP (Fig 3, lanes 5 and 6). The same is true for 1meΨTP (Fig 3, lanes 11 and 12), indicating that ΨTP and

**Figure 3.**
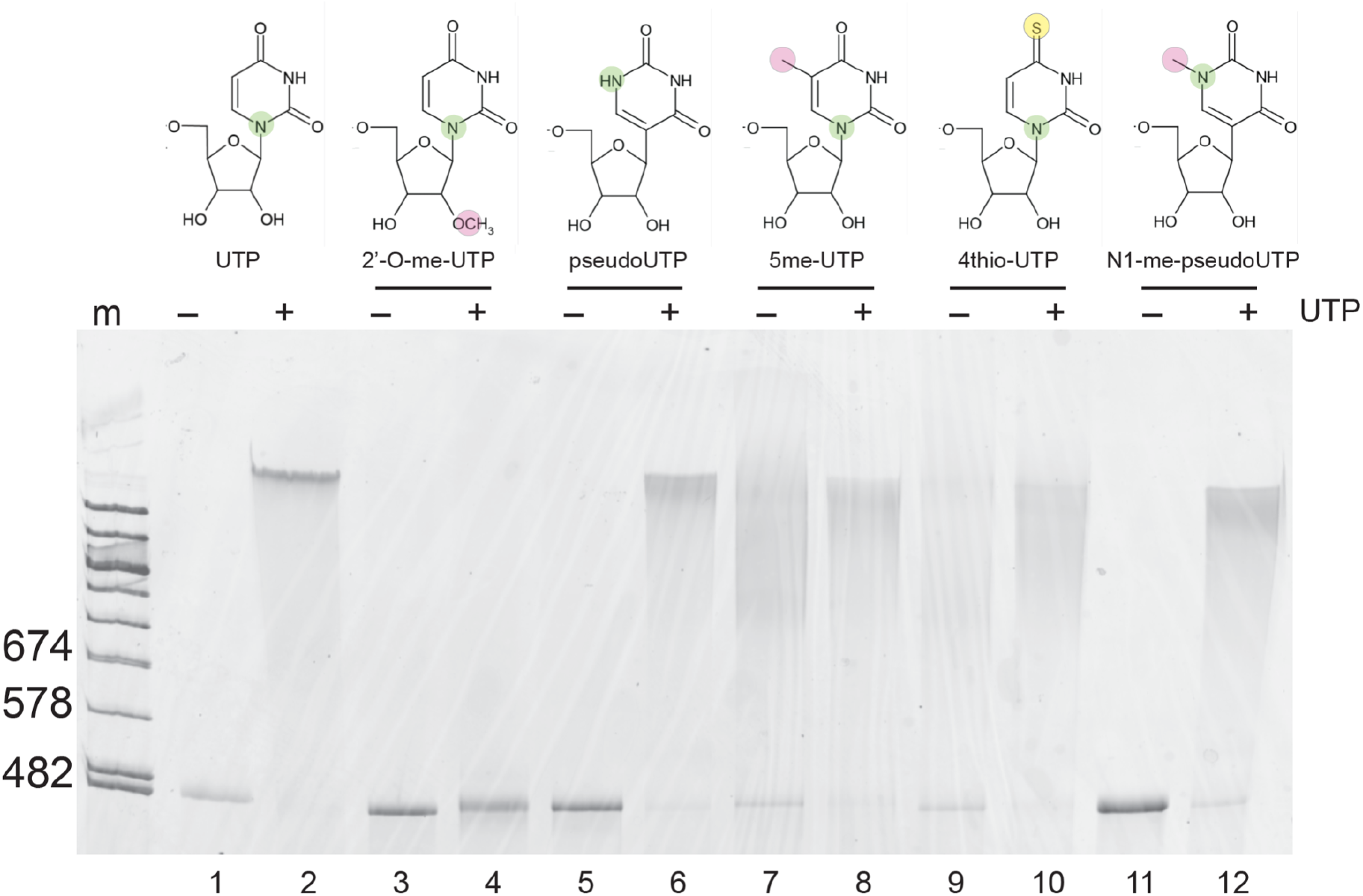
Ability of Cid1 to use modified UTP analogs for tailing. Cid1 tailing reactions were set up with the indicated analog either without (-, 1mM) or with (+, 0.5 mM each) competing unmodified UTP, using the MYL6(A+) model mRNA as a substrate. The 6% denaturing polyacrylamide gel was stained with SYBR Gold and imaged on a Typhoon Imager.

1meΨTP are inactive substrates and poor competitive inhibitors of UTP incorporation by Cid1. In contrast, Cid1 can use either 5me-UTP or 4thio-UTP, producing long tails with or without the addition of UTP (Fig 3 lanes 7, 8, 9, 10). We conclude that the ability of Cid1 to add 5me-UTP or 4thio-UTP homopolymers (or mixed modified polymers), but not ΨTP or its derivatives, has potential for additional development of nanopore RNA sequencing methods.

### Inosine tails generate a distinct signal in the nanopore during sequencing

The utility of modified homopolymers or mixed homopolymers for direct RNA sequencing depends on whether they create a distinct and recognizable signal in the nanopore. To test this we used a splint ligation method to append one to four copies of a defined synthetic inosine 15mer (I_15_) to the 3’ end of GLuc200(A+) and GLuc200(A-) RNAs (See Supplemental Methods, Suppl Fig S2B). For library preparation, we used a custom adapter with an oligo(dC) segment of 10 residues ((dC)10 adapter) that would pair with the inosine tail (Fig 4A). We purified RNA with or without a splint-ligated 30 nt poly(I) homopolymer, and with or without a poly(A) tail, prepared libraries using the appropriate adapter oligonucleotide, then sequenced the libraries on nanopores.

**Figure 4.**
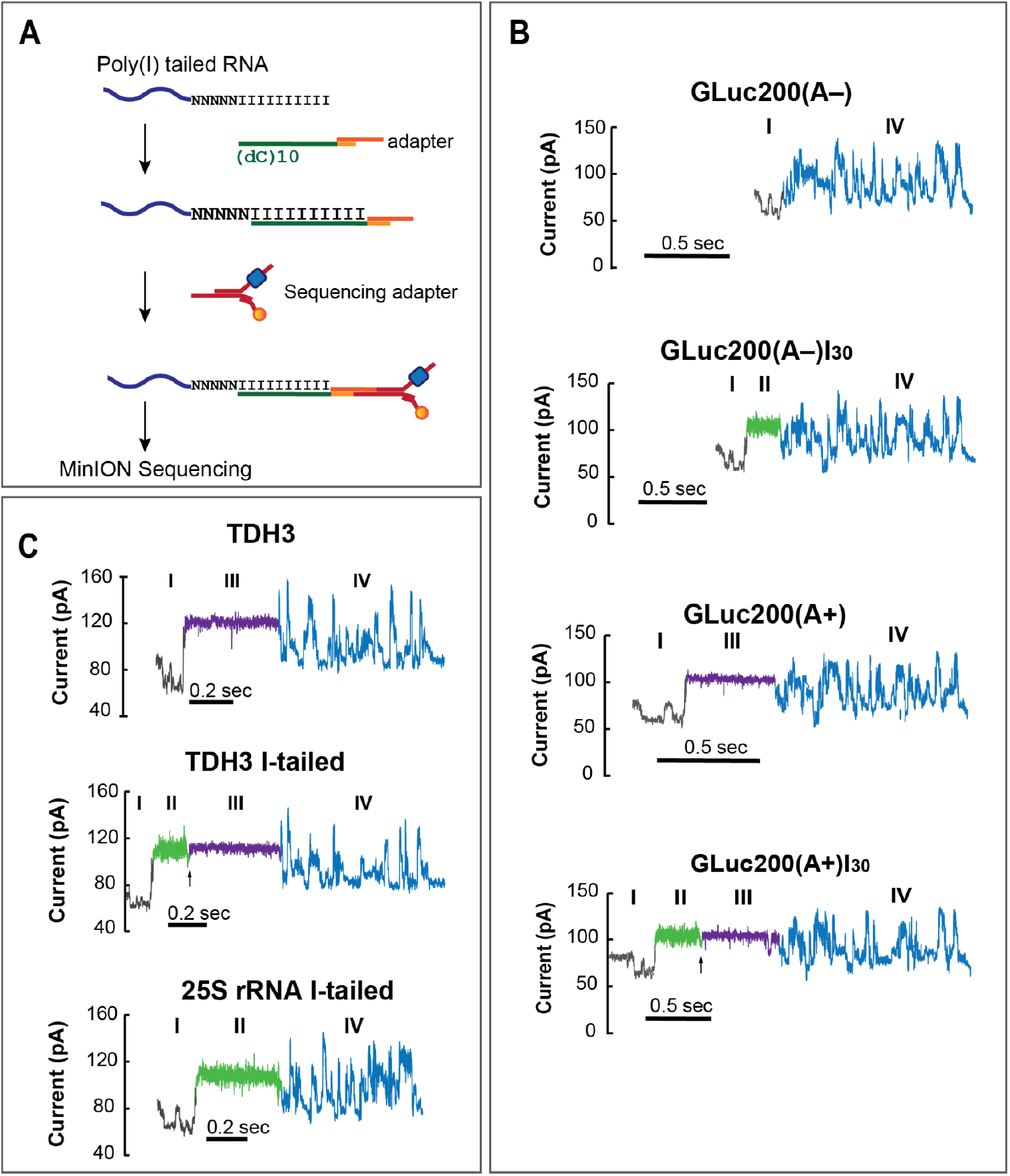
Nanopore ionic current signals for poly(I)-tailed RNA molecules. **A.** Schematic for Nanopore adaptation of poly(I)-tailed RNA molecules. This step employs a poly(dC)10 oligomer that anneals to the terminal 10 nucleotides of the added inosine tail. **B.** Ionic current traces produced by the GLuc200 model mRNA (IV, blue) with or without a poly(A) tail (II, green) and with or without a ligated 30 nt segment of inosine (III, purple). Adaptor sequence trace is shown in gray (I). RNA molecules enter the pore 3’ end first, so the trace records transit through the pore in the 3’ to 5’ direction. **C.** Ionic current traces produced by translocation of native yeast *TDH3* mRNA or 25S rRNA, with or without I-tailing by Cid1. Segments are colored as in (**B**).

A raw current trace for single representative molecules from each library is shown in (Fig 4B). Direct RNA sequencing in the ONT nanopore format threads the 3’ end of the RNA into the pore first, and the current (in picoamperes (pA), y-axis) across the pore is captured over time (x-axis) as the molecule transits the pore in the 3’ to 5’ direction. GLuc200(A-) (with no tail of any kind, but with the sequencing adapter ligated directly to its 3’ end) produces a trace showing the adapter sequence (I, gray) followed immediately by the sequence of the GLuc200 body (IV, blue, top panel). In the second panel, the splint-ligated Gluc200(A-)I_30_ molecule shows a monotonic signal at about 100 pA representing the 30 inosines (II, green) between the adapter (I, gray) and the complex sequence trace (IV, blue). By comparison, GLuc200(A+) lacking any inosines (third panel from the top) shows a monotonic signal corresponding to poly(A) (III, purple) between the adapter and the complex sequence, also around 100 pA. The bottom panel shows a molecule of Gluc200(A+)I_30_ in which the inosine tail (II, green) was splint ligated to the 3’ end of the poly(A) tail (III, purple). Although the mean ionic current associated with poly(I) and poly(A) homopolymers is similar, the ionic current noise around this mean is substantially different, with a variance of 15.1 ± 6.55 pA for poly(I) compared to 4.3 ± 3.49 pA for poly(A) (Fig 4B). In addition, there is a distinctive drop in the current amplitude (vertical arrows, Fig 4B,C) during the transition from the poly(I) to the poly(A) homopolymeric segments. We conclude that I-tails produce a distinct signal in the pore that can be distinguished from native RNA, and may be useful in distinguishing signals of native sequences from the nucleotides added for library capture. A similar current trace has been reported using I-tailing with poly(A) polymerase (Drexler et al. 2021).

### Applying the Cid1 inosine tailing reaction for capture of complex native RNA samples

Although PAP was able to add short heterogenous I-tails to poly(A)+ RNAs, the uniform +50 I-tails added by Cid1 (Fig 1) seemed better for generating representative libraries for direct RNA sequencing. Neither enzyme seemed optimal for adding inosine distributivity to all poly(A)- RNAs in a sample as a fraction of the input material remains unreacted. Since Cid1 appeared to add I-tails to poly(A)- molecules more readily than PAP (Fig 1, Fig S1), we chose to focus on Cid1. To optimize the Cid1 reaction, we incubated MYL6(A+) with Cid1 and ITP under different conditions, and measured production of the ~50 nt I-tailed product (see Methods). Under optimal conditions, Cid1 reacted immediately after addition to the reaction mixture (“0” time sample comes from adding enzyme to the reaction at 4°C, mixing and within seconds taking an aliquot to EDTA), and was complete by 40 minutes (Suppl Fig S2C). A small volume reaction with 1.2-2.4 pmol of RNA ends and 2 units of Cid1 from NEB is sufficient to convert nearly all input poly(A)+ molecules to the +50 form (Suppl Fig S2D). This is an amount of RNA that can be used as input for the commercially available Oxford Nanopore Direct RNA library protocol.

To test the use of Cid1 I-tailing for complex natural RNA samples, we I-tailed poly(A)+ RNA from *Saccharomyces cerevisiae*, prepared libraries using the oligoC adapter (Fig 4A), comparing them to standard poly(A)+ libraries made using the oligo(dT) adapter in the ONT kit (See Supplemental Tables S1 and S2 for a summary). Among the reads obtained from these libraries are examples of *TDH3* mRNA and 25S rRNA (Fig 4C, note: poly(A) selection does not completely remove rRNA, see below). As observed for the synthetic mRNA, a native yeast *TDH3* mRNA from the standard library shows the monotonic signal originating from the native poly(A) tail (top panel, III, purple) followed by the complex trace from the mRNA body (IV, blue). The second panel (Fig 4C) shows a *TDH3* mRNA that was I-tailed, and as for the synthetic mRNA (Fig 4B, bottom panel), the poly(I) signal (II, green) has the same mean current amplitude but broader variation than the poly(A) signal (III, purple), and shows the characteristic drop in current amplitude (arrow) at the transition from poly(A) to poly(I). The 25S rRNA does not have a native poly(A) tail, however I-tailing allows its capture during library preparation and sequencing. As expected, only the broad variance I-tail current trace is observed (Fig 4C, bottom panel, II, green), after which follows the complex trace formed by the 25S rRNA sequence. We conclude that I-tailing can be used to capture and analyze both poly(A)+ and poly(A)- RNA molecules from a complex natural sample, and that the variance or width of current variation observed allows poly(I) to be distinguished from poly(A), at least visually in these traces.

### Detection and quantification of poly(A)+ mRNAs using I-tailing

The ability of I-tailing to capture both poly(A)+ and poly(A)- RNAs from the same biological sample potentially extends analysis of the transcriptome beyond mRNA (Fig 4). However, to determine whether poly(I) tailing captures the mRNA population as faithfully as the direct poly(A) method, we compared multiple libraries of different preparations of yeast poly(A)+ RNA using either the commercial poly(A) capture method or the Cid1 I-tailing poly(I) capture method. Reads were mapped to the genome and counted for each gene, and coverage for each gene was compared (Supplemental Fig S3). When direct poly(A)-capture libraries are compared to I-tailed and poly(I)-capture libraries made using poly(A)+ mRNA, the efficient capture of residual rRNA reads in the I-tailed libraries distorts the parametric similarity between the libraries, but leaves a strong rank order (nonparametric) correlation intact (Fig S3).

To estimate similarity in mRNA expression measurements across methods, we filtered out rRNA reads and adjusted coverage to reads per million minus rRNA (RPM-rRNA, Fig 5A).

**Figure 5.**
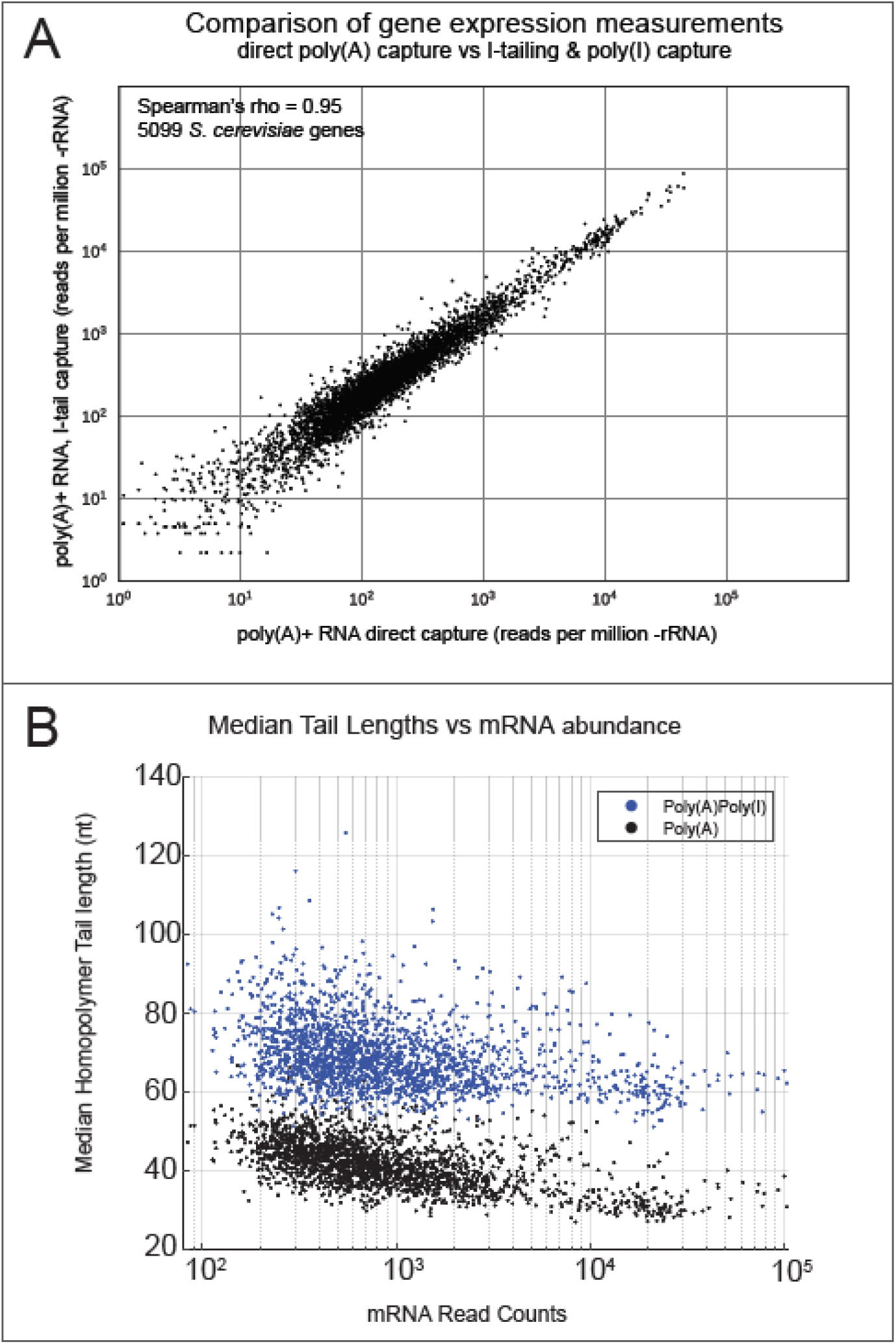
Quantitative evaluation of yeast mRNA detection after I-tailing. **A.** Strongly correlated abundance measurements for >5000 yeast mRNAs using either standard poly(A)+ capture or I-tailing by Cid1 and capture with the oligoC adapter. **B.** Estimation of median tail lengths by Nanopolish of poly(A)+ mRNAs using direct capture of poly(A)+ with the standard oligo(dT) adapter (black) or after tailing and capture with the oligoC adapter (blue). The version of Nanopolish used here has only been trained to distinguish and measure poly(A) tails (see text).

Comparison of expression of >5000 genes shows that the two methods provide highly similar measurements across the yeast poly(A)+ transcriptome (Fig 5, Pearson’s r = 0.95, Spearman’s ϱ = 0.95). Additional comparisons are shown in Fig S3 and indicate that technical replicates using the poly(I)-tailing method may be slightly more noisy than those using the commercial poly(A)-capture method, as expected for a protocol with an additional step (I-tailing). In addition, the detection of contaminating poly(A)- RNAs (not counting rRNAs which are filtered out) by the I-tailing method suggests that broader capture of snRNAs, snoRNAs, and other non-polyadenylated RNAs may explain some of the differences (see below). Overall, the two methods seem equally fit for the measurement of poly(A)+ mRNA from complex biological samples, but the I-tailing method captures additional RNAs that are not polyadenylated.

### Detecting the homopolymer extension on I-tailed poly(A)+ mRNAs using nanopores

Biochemical analysis indicates that a model poly(A)+ mRNA receives ~50 new residues during I-tailing (Fig 2B). To determine if Cid1 adds a uniform length I-tail onto different poly(A)+ mRNA molecules in a complex sample, we analyzed poly(I)-tailed total yeast poly(A) mRNA nanopore reads using Nanopolish, a software package for estimating poly(A) tail lengths (Version 0.10.2, Loman et al. 2015). Because Nanopolish is not trained to distinguish between poly(I) and poly(A), it calls the linked poly(I)-poly(A) signal as a single homopolymer segment (Fig 5B, blue). We plotted the mRNAs by coverage (x-axis) and Nanopolish median tail length estimate (y-axis) from a standard poly(A) capture library (black) and a poly(I)-tailed library (blue) that gave comparable numbers of total reads (Fig 5B, Supplemental Table S3). The majority of the control poly(A) homopolymer length estimates are in the 30-50 nt range as expected for yeast mRNAs (Sachs and Davis 1989), with the more abundant mRNAs having slightly shorter poly(A) tails on average than less abundant mRNAs (Lima et al. 2017). For the poly(I)-tailed poly(A)+ mRNAs, Nanopolish median tail length (per gene) estimates range from 60-80 nucleotides with an average of 70 nucleotides (Fig 5B). By subtraction of the median poly(A) alone tail estimates, this suggests a poly(I) tract of about 28-30 nt, well below the 50 nt determined by biochemical analysis (Fig 2). Nanopolish estimation of homopolymer lengths is at least partially dependent on the rate at which the polymer transits the pore, suggesting that perhaps poly(I) transits more quickly than poly(A). Future software development may allow computational discrimination of poly(I) segments from poly(A) segments within each homopolymer tails as well as more accurate estimate of I-tail lengths. The strong agreement in gene expression measurements made with or without I-tailing indicates that Cid1 uniformly adds about the same length I-tail to each poly(A)+ mRNA, with the exception of very short mRNAs as noted above (Figs 1, S1).

### Capture and measurement of non-polyadenylated RNAs by I-tailing

To examine the ability of Cid1 to I-tail non-polyadenylated RNAs, we incubated total and rRNA-depleted yeast RNA samples with Cid1 and ITP and (Fig 6). After Cid1 I-tailing, a slight shift up in migration of the 18S and 25S rRNAs is observed (Fig 6A, arrows). Because purified 5.8S rRNA acquires a heterogenous I-tail that can be quite long (Fig 2D), 5.8S rRNA no longer appears as a sharp band on the gel (Fig 6A). There is little if any distinguishable change in 5S rRNA or the tRNAs, suggesting they may fail to bind the enzyme or their 3’ ends may be buried or otherwise blocked from entering the enzyme active site. These results reinforce observations using model RNAs (Fig 1) that different non-polyadenylated RNAs may have different Cid1 tailing efficiencies.

**Figure 6.**
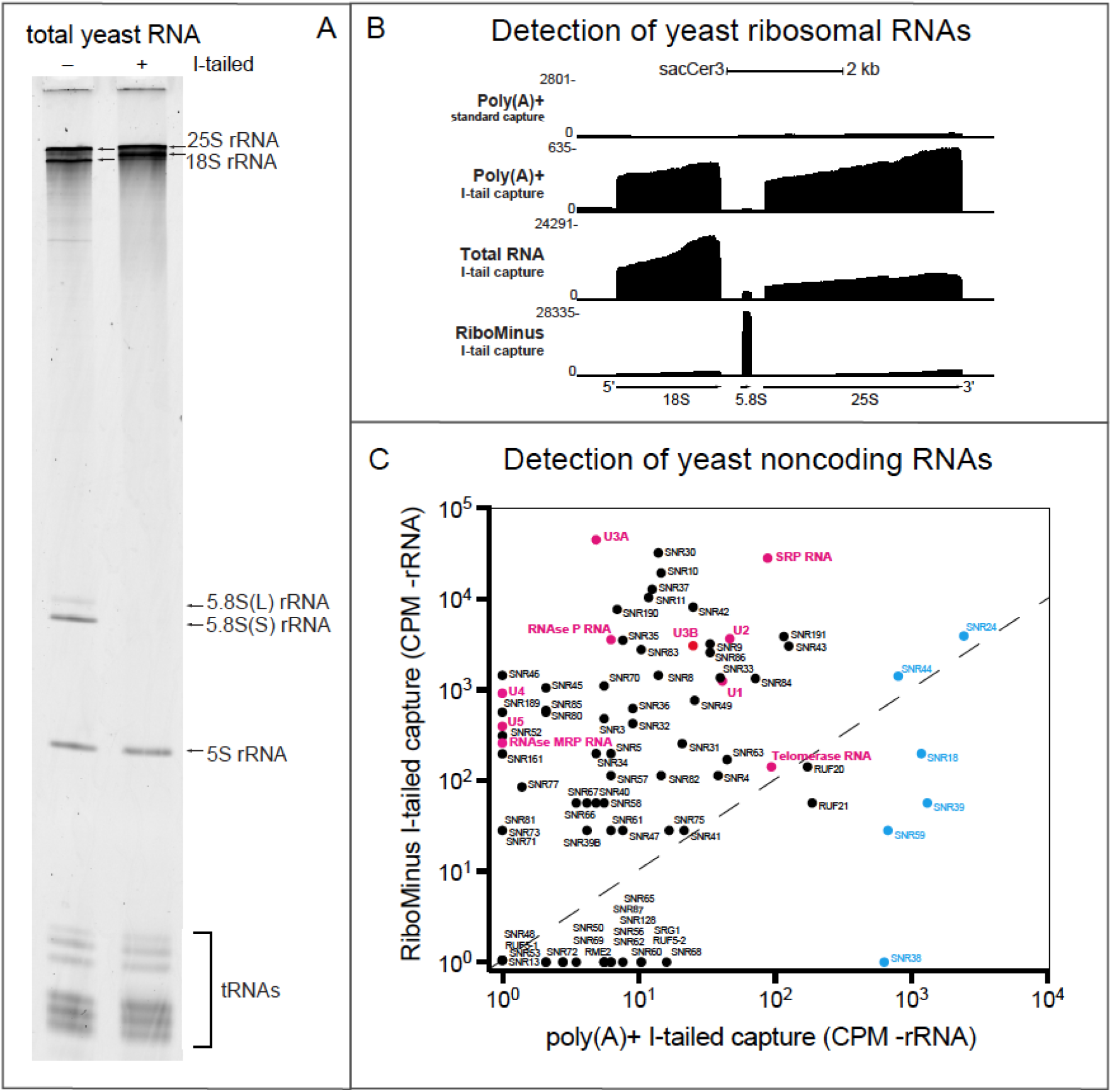
Detection of non-polyadenylated RNAs. **A.** I-tailing of total yeast RNA. Yeast RNA treated with (+) or without (-) Cid1 and ITP for I-tailing were run on a 6% denaturing polyacrylamide gel and stained with SYBR Gold. Slight shift up of 18 and 25S rRNAs indicates addition of poly(I). Disappearance of 5.8S rRNA is consistent with its acquisition of a long heterogeneous tail (See Fig 2). **B.** genome browser view of the rRNA genes of *S. cerevisiae* showing detection of rRNA by different library preparation methods. Top line is poly(A) selected RNA using the direct oligoT capture method, second line is poly(A) selected and I-tailed RNA captured with the oligoC adapter, third line is total RNA I-tailed and captured with the oligoC adapter, fourth line is Ribominus rRNA depleted total RNA I-tailed and captured with the oligoC adapter. **C.** Scatter plot showing robust detection of non-polyadenylated noncoding RNAs by I-tailing and capture with the oligoC adapter.

To determine the extent to which I-tailing by Cid1 is useful for detecting and measuring non-poly(A) RNA, we sequenced rRNA from libraries made using different RNA sources by both the standard poly(A) capture method and the I-tailing method. Libraries made from either poly(A) selected RNA (rRNA is a contaminant in poly(A) selected RNA), total RNA, or total RNA subjected to rRNA depletion by the RiboMinus kit were sequenced on nanopores and mapped to the *S. cerevisiae* rDNA locus (Fig 6B). As expected, the standard ONT poly(A) capture libraries have less than 0.1% of reads aligning to the rRNA genes (Supplemental Table S2). By comparison, when poly(A)-selected RNA is I-tailed and captured using the oligoC adapter, residual non-polyadenylated RNAs contaminating the poly(A) selected sample is detected with ~4-8% of the reads comprising rRNA (Supplemental Table S2). In total RNA with no depletion step, I-tailed rRNAs accounted for 73% of the reads (Supplemental Table S2), suggesting that sensitivity of detection of other non-polyadenylated RNAs would be improved by rRNA depletion. RiboMinus treatment successfully depleted 18S and 25S rRNA, but not 5.8S rRNA (Fig 6B), which still comprised 41% of the reads from this sample (Supplemental Table S2). Since the much shorter 5.8S rRNA takes far less time than 18 or 25S rRNA to go through the pore, Ribominus rRNA depletion allowed detection of a diverse set of non-polyadenylated RNAs including snoRNAs (Fig 6C, black circles), the yeast homologs of telomerase RNA, RNAseP RNA, U3 snoRNA, and all the spliceosomal snRNAs except U6 (which has a 3’ phosphate that is not a substrate for I-tailing, Lund and Dahlberg 1992), and others (Fig 6C, magenta circles). Reads mapping to the intronic snoRNA genomic coordinates (blue circles) are efficiently captured in poly(A)+ samples as part of the pre-mRNA precursors and processing intermediates within which they reside. Our ability to detect noncoding RNAs is likely a product of both their abundance in the sample and in the accessibility of their 3’ ends to the enzyme.

### Capturing nascent transcript structure by Cid1 I-tailing of chromatin-associated RNA

There is growing interest in methods that can capture and directly sequence nascent RNAs, enabling RNA modification and processing to be detected co-transcriptionally (Wissink et al. 2019; Hsiao et al. 2018; Incarnato et al. 2017). We investigated the suitability of I-tailiing for detecting nascent RNA transcripts by enriching for nascent RNAs and isolating chromatin from yeast strain CKY2647 (gift from Dr. Craig Kaplan). CKY2647 has a C-terminal AviTag on the RNAPII *RPB3* subunit and constitutively expresses the biotin ligase BirA, which biotinylates the AviTag in vivo (Fairhead and Howarth 2015). Chromatin preparations were incubated with streptavidin beads to capture biotin-tagged RNAPII along with nascent transcripts, and then recovered by phenol-chlorofom extraction (Fig 7A).

**Figure 7.**
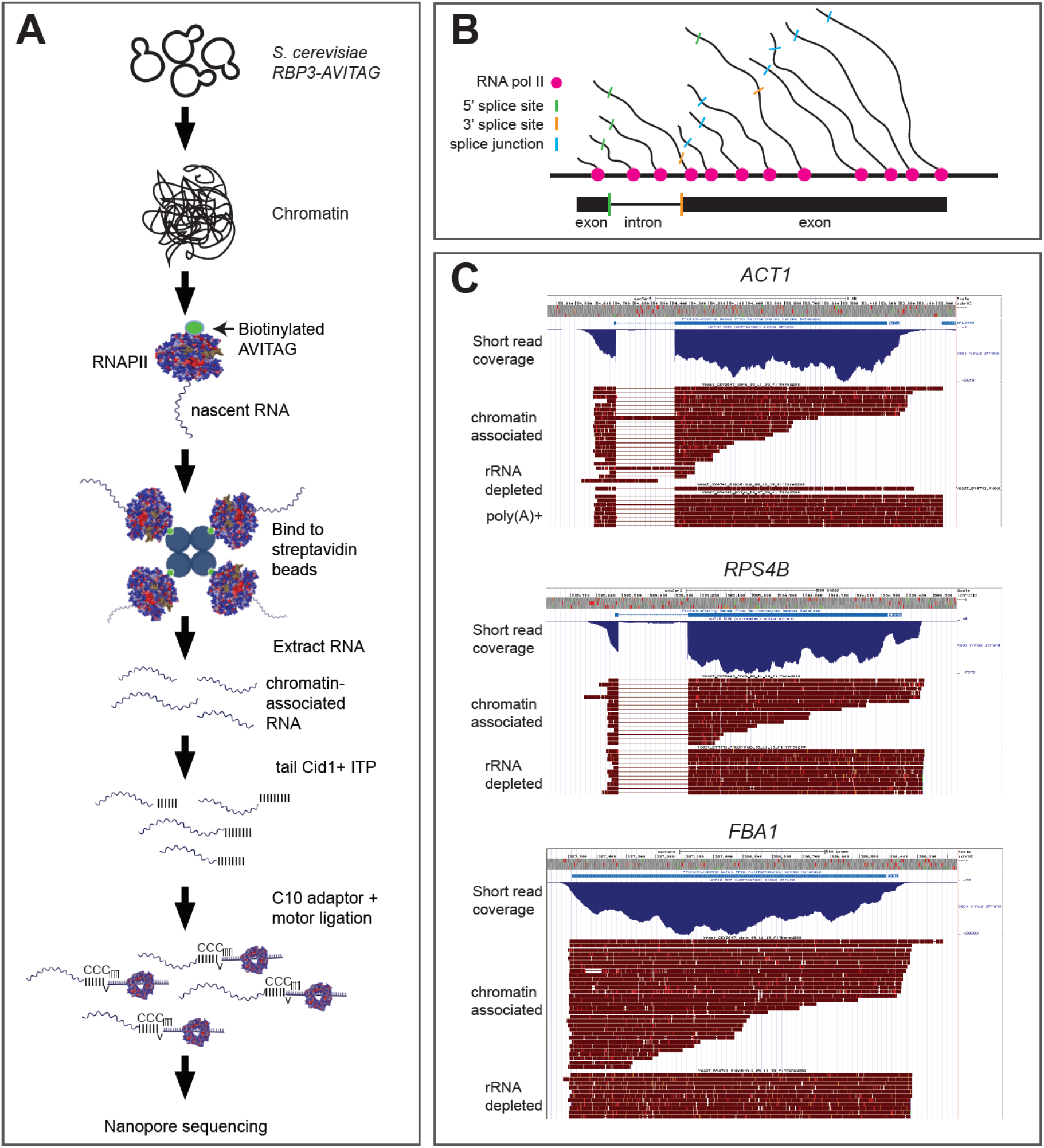
Capture and analysis of RNA polymerase II (RNAPII) associated nascent transcripts by Cid1 I-tailing. **A.** Protocol for isolating RNAPII-associated transcripts from chromatin (see methods). **B.** A model for co-transcriptional splicing. Introns bounded by 5’ splice sites (green) and 3’ splice sites (gold) are removed creating exon-exon junctions (blue) in nascent transcripts as RNAPII (magenta) moves down the gene. **C.** genome browser views of nanopore reads mapping to ACT1 (top), RPS4B (middle), and FBA1 (bottom). The 3’ ends of reads from nascent RNAs are distributed along the gene, and splicing usually occurs before RNAPII has moved 100 nt beyond the 3’ splice site.

After purifying the RNA from this sample, we I-tailed, sequenced, and mapped it to the yeast genome, inferring that the location of the 3’ end of the transcript would identify the position of RNAPII on the gene (Fig 7B, Churchman and Weissman 2011). To analyze transcripts that initiated at the promoter, we filtered for reads whose 5’ ends mapped within 200 bp of the annotated start codon and aligned these to the genome, comparing them to similarly filtered transcripts from the rRNA depleted I-tailed library (*RPS4B, FBA1*) or the poly(A) selected I-tailed library (*ACT1*, Fig 7C). Whereas >90% of the transcripts found in rRNA depleted or poly(A) selected total RNA have 3’ ends that map together near the end of the transcription unit (as judged by short read coverage, blue tracks, Fig 7C), the 3’ ends of transcripts from the chromatin-associated RNA were distributed along the gene as expected for nascent transcripts. For the intron-containing genes *ACT1* and *RPS4B*, most of the nascent transcripts have had their introns removed while RNAPII is still engaged, consistent with co-transcriptional splicing. In agreement with other work (Carrillo Oesterreich et al. 2016), a substantial fraction of yeast nascent transcripts are spliced before RNAPII moves more than 100 nt from the 3’ splice site (Fig 7C). We conclude that following preparation of chromatin-associated RNA, I-tailing by Cid1 is suitable for detection and analysis of processing of nascent transcripts by nanopore direct RNA sequencing.

## DISCUSSION

We set out to test the ability of 3’ nucleotidyl transferases to incorporate modified nucleotides in order to explore potential applications for nanopore direct RNA sequencing. Polynucleotide phosphorylase, useful for base non-specific addition of rNDPs to 3’ ends (Grunberg-Manago et al. 1956; Grunberg-Manago 1989) was initially tested and ruled unsuitable due to 3’ trimming by the competing reverse 3’ phosphorolytic activity (for example, see Unciuleac et al. 2021). Upon testing Cid1 poly(U) polymerase, new activities of Cid1 were revealed (Figs 1-3), in particular the unexpected synthesis of inosine homopolymers (Fig 2). We exploited this new activity to capture synthetic and natural transcripts from complex samples for nanopore sequencing (Figs 4-7). We showed that I-tailing by Cid1 does not distort measurement of poly(A)+ mRNAs (Fig 5), while simultaneously enabling robust detection of essential non-polyadenylated RNAs such as snRNAs, telomerase, and RNAse P RNA (Fig 6). Composition of RNAs isolated from chromatin, such as partially processed nascent transcripts, can also be analyzed (Fig 7).

### New activities of Cid1

Our examination of the ability of Cid1 to add modified nucleotides to the 3’ ends of RNA was motivated by the limitations of the standard nanopore direct RNA sequencing protocol. The standard protocol targets the natural poly(A) tail present on most mRNAs (Garalde et al. 2018), however many target RNAs, such as nascent RNAs, mature snRNAs, snoRNAs, rRNAs and many other noncoding RNAs, lack a poly(A) tail. Enzymatic addition of poly(A) to capture non-polyadenylated RNAs (Wongsurawat et al. 2019; Drexler et al. 2021) complicates the situation by adding information that cannot be discerned from that present on natural RNAs undergoing decay or other post-transcriptional events that involve addition of short poly(A) tracts (Tudek et al. 2018), or other 3’-nucleotidylation events (Zigáčková and Vaňáčová 2018; Liudkovska and Dziembowski 2021). ATP analog 6-bioATP (N6-[(6-amino)hexyl]-amino-ATP-biotin is a substrate for Cid1 (Moritz and Wahle 2014), but evidence for other non-canonical nucleotides was scarce, and we were uncertain whether such a large modification would fit through the pore. Our results indicate that addition of modified nucleotides is a feasible alternative to the standard method and that distinct current signatures allow their discrimination from native nucleotides.

The structure and function of Cid1 has been studied in detail (Yates et al. 2012, 2015; Munoz-Tello et al. 2012, 2014; Lunde et al. 2012; Rissland et al. 2007), however it remains unclear how ITP is recognized by the enzyme. Crystal structures of Cid1 bound to each of the four standard rNTPs (Lunde et al. 2012), show that while U, A, and CTP are bound in the *anti* orientation, GTP is bound in the *syn* orientation, possibly because the guanosine 2-amino group fits poorly into the binding pocket otherwise. Inosine lacks the 2-amino group of guanosine, suggesting it may be able to bind in the *anti* conformation, similar to ATP or UTP (Lunde et al. 2012). In this orientation, the N1 and O6 of inosine might interact with enzyme functional groups that normally interact with N3 and O4 of its preferred substrate UTP (Lunde et al. 2012; Yates et al. 2012; Munoz-Tello et al. 2012). Additional studies will be needed to explain how inosine is recognized by Cid1.

Our survey of UTP analog use also revealed interesting new properties of the Cid1 enzyme. First, 2’-O-methyl-UTP, strongly inhibits UTP incorporation while supporting extension of the RNA substrate by one or a few nucleotides at most (Fig 3), suggesting it can be incorporated but is chain terminating. Second, the enzyme uses 5meUTP (aka ribothymidine triphosphate) but not ΨTP or 1meΨTP (Fig 3). This is surprising because the methyl group on the C5 position of U in 5meUTP occupies the same space as the methylated N1 in 1meΨTP (Fig 3, note that atoms at positions 5 and 1 are flipped during conversion of UTP to ΨTP, Spedaliere et al. 2004). It is hard to imagine how this difference would affect binding or catalysis given the available crystal structures (Munoz-Tello et al. 2014; Lunde et al. 2012; Yates et al. 2012; Munoz-Tello et al. 2012). It seems unlikely that the structural basis for discrimination against ΨTP analogs lies in the difference between the glycosidic bond (N1 to C1’) in UTP and 5meUTP as compared to the C-C bond (C5 to C1’) joining base to sugar in ΨTP and 1meΨTP, but other explanations are not evident and will require future experiments.

Addition of inosine tracts to poly(A)+ mRNA abruptly stalls after addition of about 50 residues (Fig 1C, 2B, 5B). Although we have not systematically determined them, the requirements for this strong stop appear to include a specific length of poly(A) greater than 24, since we do not observe this stop in the oligo A24 reaction (Fig 1A). We have not tested a model mRNA with an A-tail of 24 so it is also possible that additional RNA 5’ of the A tract may be required to produce the +50 stop. A 200 nt model mRNA with a 44 nt poly(A) tail was a substrate for A and U addition, but tailed poorly with I, showing low amounts of long I-tails with no accumulation of +50 nt product (GLuc200A44, Fig S1A), suggesting that features in the mRNA body may contribute to the quality of binding to Cid1. Most yeast mRNAs have >24 A residues and the high degree of quantitative correspondence between I-tailed libraries with the standard poly(A) capture libraries (Fig 5) indicates that few if any natural yeast mRNAs behave like the GLuc200A44 substrate. What creates the strong stop is unknown, however I-A base “wobble” base pairing could produce an extended duplex or other structure that might prevent further reaction once the I-tract reaches 50 residues.

RNAs that lack poly(A) tails acquire a heterogeneous length I-tail that ranges from a few to many hundreds of residues long (Fig 1A, C Fig 2C, D). The per molecule efficiency of I-tailing on RNAs lacking poly(A) tails is variable, and the product distribution suggests that initially the reaction is distributive and inefficient until some molecules acquire a short I-tail after which the reaction becomes more processive on that class of molecules. An example of an efficient substrate is 5.8S rRNA (most molecules immediately acquire heterogenous tails), whereas 5S rRNA is barely if at all recognized (Fig 6). The minimum size for initial substrate recognition by Cid1 is at least 15 residues (Rissland et al. 2007), and we can easily detect 5.8S rRNA (~120 nt) and several snoRNAs that are less than 100 nt. The characteristic tailing efficiency of each individual poly(A)- RNA will influence its sensitivity of detection. We have not examined the effect of the length of the I-tail on capture efficiency during library preparation or sequencing.

### Applications and limitations of the current method and potential for future development

We have demonstrated the utility of Cid1 I-tailing for capture, library preparation, and direct RNA sequencing on nanopores. One particular advance from this effort is the ability to detect many important non-polyadenylated structural RNAs in the transcriptome (Fig 6). Because the 3’ ends of many of these RNAs are buried in a structure that helps stabilize them, their Cid1 I-tailing efficiencies will be idiosyncratic, and thus their representation in libraries may not match their abundance in the sample. This limitation is present for any method that requires access to the RNA 3’ end, and it means determination of absolute numbers of molecules or quantification of two such RNAs relative to each other within the same sample may not be possible. Nonetheless, the relative change in the same RNA measured in two samples may be determined, provided the treatment does not alter the 3’ end of the target RNA. Thus the method will capture changes in both the amount of an RNA and in the structure of its 3’ end. The ends of most nascent RNA transcripts are unlikely to be highly structured as they will be distributed along all positions of the transcription unit, and thus representation should be relatively even across the gene (Fig 7). Thus this method enables detection of a large and complex class of RNA that are missed by the standard poly(A) capture method. At the same time we show that I-tailing quantitatively recovers poly(A)+ mRNAs equivalently to the standard method, allowing both poly(A)+ and poly(A)- RNAs to be captured in the same sample.

Our poly(I) sequencing libraries typically have 10-30% the throughput compared to standard ONT poly(A) sequencing (Table S2), potentially a concern for a method that is already limited by throughput. One reason could be that the long I-tails on some RNAs sequester the C10 adapter, preventing its ligation to other molecules, thus reducing the efficiency of ligation of the motor protein adapter, and reducing the depth of the library. Another possibility is that very long poly(I) tails may be interpreted as stalls in the nanopore by the software used to run the sequencer, MinKNOW. Any monotonic state in the ionic current that persists for 5 seconds or longer is considered a stall, and MinKNOW reverses the voltage across the nanopore to eject any strand trapped in the nanopore. During DNA sequencing in nanopores, molecules that are ejected are unlikely to be sequenced since the motor protein has already migrated across the molecule (Loose et al. 2016). Further development to control the length of the I-tails may address both of these possible limitations to throughput.

I-tailing provides a distinct electronic signature (Fig 4), potentially allowing RNAs from an I-tailed sample to be distinguished from non-I-tailed molecules in a sequencing experiment where two different libraries have been mixed. Currently the ability to discriminate I-tails from A-tails from I+A-tails is insufficiently accurate to deconvolute read sample origin (unpublished, see also Drexler et al. 2021). In addition several modified UTP analogs (Fig 3), as well as at least one N6 modified ATP analog (Moritz and Wahle 2014) can be incorporated by Cid1. No doubt Cid1 can create mixed polymers from combinations of base modified precursors and these may very well have distinct electronic signatures that will allow a tailing step to double as a barcoding step for direct RNA sequencing in nanopores. Additional 3’-nucleotidyl transferases besides Cid1 may have additional capabilities that have yet to be exploited (e. g. Preston et al. 2019).

## MATERIALS AND METHODS

### *In vitro* transcription template generation

DNA templates for MYL6(A+) and MYL6(A-) transcripts were generated using linearized pUC13-MYL6 plasmid. pUC13-MYL6 was digested with BbsI (NEB: R0539S, cleaves after the poly(A) sequence in the template), or BsmI (NEB: R0134S, cleaves before the poly(A) sequence in the template) for MYL6(A+) or MYL6(A-) respectively (see Supplemental Methods for sequence of the MYL6 template. Gluc200 and Gluc200A44 templates were generated using PCR amplification of GLuc of the first 200 nt at the 5’end of pCMV-GLuc 2 Control Plasmid (NEB: https://www.neb.com/tools-and-resources/interactive-tools/dna-sequences-and-maps-tool). The sequence was targeted using a forward primer (5’-TCGAAATTAATACGACTCACTATAGGGAGACCCAA) containing a T7 promotor region, and a reverse primer that terminates the PCR product at the 3’ end of the truncated GLuc sequence (5’-GCGGCAGCTTCTTGCC) or templates the addition of a 40 nt 3’ poly(A) tail for GLuc200 and GLuc200A44 (5’-TTTTTTTTTTTTTTTTTTTTTTTTTTTTTTTTTTTTTTTTTTTTGCGGCAGCTTCTTGCC) respectively. Amplification was obtained using Platinum^®^ Taq DNA Polymerase High Fidelity (Invitrogen) in reactions with 1x High Fidelity PCR buffer, 0.01ug/ul of plasmid, 0.4mM dNTP Mix, 0.4uM Forward Primer, 0.4uM Reverse primer, 2mM MgSO_4_, 1unit Platinum^®^ Taq DNA Polymerase High Fidelity. PCR cycles were: 94C 30s, [94C 10s, 58C 15s, 65C 45s] 22 cycles, then 65C 10m, 4C hold. DNA In vitro transcription templates were purified using NucleoSpin^®^ Gel and PCR Clean-up (Macherey-Nagel) following the manufacturer’s instructions.

### T7 *in vitro* transcription

Templates (described above) were transcribed using the MEGAscript™ T7 Transcription Kit (Invitrogen). Reaction products were separated using 6% 8M urea polyacrylamide gel electrophoresis, identified by staining with ethidium bromide, and excised. Gel slices were rotated at 4°C overnight in RNA Elution Buffer (0.3M NaOAc pH 5.2, 0.2% SDS, 1mM EDTA, 10ug/mL proteinase K). Eluted product was purified using 25:24:1 phenol:chloroform:isoamyl alcohol and ethanol precipitation.

### RNA extraction and initial sample preparation

Total RNA from *S. cerevisiae* BY4741 cells was extracted by a hot phenol method as described previously (Ares 2012). Poly(A) RNA was selected using NEXTflex Poly(A) beads (BIOO Scientific Cat#NOVA-512980) according to the manufacturer’s instructions. Ribosomal RNA depletion of total RNA employed RiboMinus Transcriptome kits and Concentration Modules (Invitrogen #K155001, K155003) and was done according to the manufacturer’s instructions.

### Polynucleotide tailing with *S. pombe* Cid1 Poly(U) Polymerase

To test for tailing ^32^P-labeled A24 as a substrate using residues (ATP, CTP, GTP, UTP or ITP) a reaction containing labeled A24 RNA, 0.5mM NTP, 50mM NaCl, 10mM MgCl_2_, 10 mM Tris-Cl pH 7.9, 1mM DTT, 500 ug/ml Bovine Serum Albumin (BSA), and 0.6 units of NEB poly(U) polymerase in a final volume of 5ul was incubated at 37°C for 30 minutes. The standard reaction for adding homopolymers to the 3’ends of model mRNAs and yeast RNA preparations using Cid1 poly(U) polymerase is as follows: RNA (various amounts) in 0.1mM EDTA in volumes up to 2.95uL is denatured at 95°C for 2 minutes then placed on ice for 2 minutes. The RNA is added to a reaction containing 4mM NTP (ATP, CTP, GTP, UTP or ITP) 50mM NaCl, 13.5mM MgCl_2_, 10 mM Tris-Cl pH 7.9, 1mM DTT, 500 ug/ml BSA, and 0.6 units of NEB poly(U) polymerase in a final volume of 7.5ul and incubated at 37°C for 1 hour. To test for incorporation of modified U residues, 200fmol MYL6(A+) RNA was incubated with 1mM concentration of modified UTP analog or a combination of 0.5mM modified UTP analog and 0.5mM UTP following the standard poly(U) polymerase reaction.

Inosine triphosphate was purchased from Sigma. Modified UTP analogs were purchased from TriLink, Inc.

### Polynucleotide tailing with *S. cerevisiae* Poly(A) Polymerase

To add a homopolymer to the 3’ ends of RNAs using Poly(A) Polymerase, Yeast (ThermoScientific 74225Y/Z) a reaction containing (1x Poly(A) Polymerase reaction buffer, 200fmol RNA, 0.5mM NTP) is incubated at 37°C for 30 minutes, then two volumes of Gel Loading Buffer II (Invitrogen: AM8546G) are immediately added to stop the reaction. Reaction products were separated on 15%, 8%, or 6% denaturing polyacrylamide gels depending on their sizes.

### pCp labeling

T7 transcripts and their I-tailed counterparts were labeled with pCp [5’-32P] Cytidine 3’, 5’ bis(phosphate) 3000 Ci/mmol, 10mCi/ml, using NEB T4 RNA Ligase 1, 1X reaction buffer, 0.15mM ATP, 10% DMSO, 20 uCi ^32^P-pCp, 6pmol of RNA, and 333 units RNA Ligase 1 (NEB M0204S) with incubation at 16°C for ~16-18 hours. The products were purified by extracting with an equal volume of 25:24:1 phenol:chloroform:isoamyl alcohol and 0.3mM NaOAc pH 5.2 was added before ethanol precipitation.

### RNAse A digestion

RNA samples in 100mM NaCl, 10mM EDTA, 0.025ug/ul RNAse A (Thermo Scientific RNase A, Cat No EN0531) were incubated at 37°C for 15 minutes, then immediately vortexed with equal amounts of 25:24:1 phenol:chloroform:isoamyl alcohol. The aqueous phase was made 0.3 M NaOAc pH 5.2 precipitated with ethanol, rinsed with 70% ethanol, briefly dried and then suspended in formamide and dyes for electrophoresis.

### Preparation of RNAPII-associated RNA

#### Growth of yeast

*S. cerevisiae* CKY2647 (*MAT**a** ura3-52::BirA::kanMX his3Δ200 leu2Δ met15Δ0 trp1Δ63 lys2-128δ gal10Δ56 rpb1d::CLONAT rbp3::AVITAG-TAP::KITRP1*, pRP112 RPB1 CEN URA3) is grown to an A600 of 0.5-0.8 in YEPD in 100mL cultures. Cells are harvested at 1100 x *g* centrifugation for 5 minutes at 4°C. Pellets are washed with 40ml ice-cold PBS twice, then transferred to 1.7ml Eppendorf tubes and washed with cold PBS by centrifugation at 1100x g for 5 minutes at 4°C. Supernatant is removed and pellets are snap frozen in liquid nitrogen before storing in −80C.

#### Preparation of chromatin

A hole is pierced in the bottom of a 15mL Falcon tube with a 22-gauge needle and placed inside a 50mL Falcon tube cap into which a hole has been cut just large enough to fit the 15mL tube. A strip of parafilm is wrapped around the 12mL mark on the 15mL Falcon tube for stabilization in the 50 mL Falcon tube. The 50mL Falcon tube cap containing the 15mL centrifuge is then screwed on to the 50mL Falcon tube - this assemblage is a make-shift bead filter. Working in a 4°C room, the yeast pellets are retrieved from the −80C freezer and placed on ice for 5 minutes, then resuspended in 1ml of Buffer 1 (20mM HEPES pH 8.0, 60mM KCl, 15mM NaCl, 5mM MgCl_2_, 1mM CaCl_2_, 0.8% Triton-X100, 0.25mM sucrose, with freshly added 0.5mM spermine and 2.5mM spermidine). The cells are lysed in a 2ml Eppendorf tube containing 1mL 0.5mm zirconia beads by vortexing for six cycles of 1 minute vortexing with 1 minute pauses using the Turbomix attachment on a Vortex Genie 2 (Scientific Industries Inc SKU: SI-0564) that had been pre-run at max speed for 1 minute preceding the vortexing to ensure consistent machine performance in the cold. The sample (plus beads) is transferred to the 15mL Falcon tube. For transfer of remaining lysed cells left in the 2ml tube, 1ml of Buffer 1 was added, then the tube was inverted before transferring to the assembled bead column. This process was repeated three more times, for a total of four times altogether. The assembled Falcon tube bead filter was centrifuged at 400 x g for 6 minutes at 4°C to remove the lysate from the zirconia beads. Avoiding the pellet, 2 x 750ul of the supernatant at the bottom of the 50ml Falcon tube is transferred into each of two 1.7ml Eppendorf tubes and the pellet is discarded. The samples are centrifuged at 2000 x g for 15 minutes at 4°C, the supernatant fluid is removed and the chromatin pellets are resuspended in 800ml of Buffer 1, the pairs are combined into one 1.7ml Eppendorf tube and centrifuged again at 2000 x g for 15 minutes at 4°C. Using a p1000, the chromatin pellet is aggressively resuspended in 800ul of Buffer 2 (20mM HEPES pH 7.6, 450mM NaCl, 7.5mM MgCl_2_, 20mM EDTA, 10% glycerol, 1% NP-40, 2M urea, 0.5M sucrose, with freshly added 1mM DTT and 0.2mM PMSF). The sample is vortexed for 5-10 seconds, then incubated on ice for 5 minutes. After centrifugation at 2000 x g for 15 minutes at 4°C, the pellet is again suspended in 800ul of Buffer 2. The sample is centrifuged at 2000 x g for 15 minutes at 4°C, then the pellet is finally suspended in 300ul of Buffer 3 (HEPES pH 8.0, 60mM KCl, 15mM NaCl, 5mM MgCl_2_, 1mM CaCl_2_) before adding to prepared streptavidin beads for RNPII capture. Streptavidin beads (NEB #S1420S) are prepared by taking 150ul of the bead suspension and equilibrating them in 700ul of bead buffer (0.5M NaCl. 20mM Tris pH 7.5, 1mM EDTA), and recovering them on the wall of the tube with a magnetic rack. The bead buffer wash is discarded and the beads are resuspended in 300ul of the pellet in Buffer 3 from above. The mixture is rotated at 4°C for 2 hours, and then the beads are washed three times as follows: Collect on the tube wall on a magnetic rack, remove and discard supernatant, add 500ul Buffer 3 to the beads, remove from the magnet and suspend beads. For RNA purification, the beads are collected from the final wash and resuspended in 500ul of RNA Extraction Buffer (0.3M NaOAc pH 5.3, 1mM EDTA, 1% SDS), 100ul acid phenol, and 100ul chloroform. This is vortexed vigorously and then centrifuged at 14,000 x g for 5 minutes. The aqueous phase is combined with 1.2mL of 100% ethanol in a fresh 1.7mL Eppendorf tube, mixed well and incubated at −80°C for 1 hour or overnight. Precipitated RNA is recovered by centrifugation, the pellet is rinsed with 70% ethanol and briefly air dried. To remove contaminating DNA, the pellet is resuspended in 100ul of DNAse solution (1x Turbo DNAse Buffer, 10 units TURBO DNAse (Invitrogen AM2238)) and incubated at 37°C for 30 minutes. The sample is finally purified using a RNA Clean and Concentrator-5 kit (Zymo R1013) following the manufacturer’s instructions, and eluted in 35ul of RNAse-free water.

### Library preparation for direct RNA sequencing on nanopores

Purified RNA (500-775 ng) was prepared for library construction as follows: Poly(A) enriched samples captured by their endogenous poly(A) tail using either the ONT SQK-RNA001 kit or its replacement SQK-RNA002 kit following the manufacturer’s instructions. I-tailed RNA was captured using the ONT SQK-RNA002 kit except that a custom oligoC adapter was used in place of the RTA (oligoT) adapter provided by the kit. The optional reverse transcription step was done for all libraries in this study except that Superscript IV (Thermo Fisher) was used in place of Superscript III. To assemble the custom oligoC adapter, 100 pmol of top oligo (5’-pGGCTTCTTCTTGCTCTTAGGTAGTAGGTTC-3’) and 100pmol bottom oligo (5’-CCTAAGAGCAAGAAGAAGCCCCCCCCCCCC-3’) are mixed in 10 ul of 50mM Tris pH 8, 100mM NaCl, 0.1mM EDTA and incubated in a thermocycler at 75°C, then slow cooled at a ramp rate of 0.1°C/sec to 23°C. The annealed duplex adapter is diluted to 100μl with 90μl water, and 1 μl of this 1 uM duplex was used for capture of I-tailed RNA. Libraries were sequenced on the MinION using ONT R9.4 flow cells and the standard MinKNOW protocol script RNA001 or RNA002 recommended by ONT with one exception: bulk phase raw files were collected for the first 2 hours of sequencing and then standard sequencing was restarted and commenced for ~48 hours.

### Base-calling and mapping reads to the yeast genome

We used Guppy Base-calling Software from Oxford Nanopore Technologies, Limited. Version 3.0.3+7e7b7d0 with the configuration file “rna_r9.4.1_70bps_hac.cfg” (Wick et al. 2019) was used for base-calling direct RNA traces. NanoFilt version 2.5.0 (Coster et al. 2018) was used for classification of passed reads. Reads classified as “pass” had phredscore threshold of ≥ 7 and “failed” if < 7. The alignments in Figure 7 were base called using Guppy version 4.4.2+9623c16 with the same configuration file as above, but this did not result in significant changes to mapping or read counts. Passing reads were mapped to the sacCer3 yeast genome using minimap2 version 2.16-r922 (Coster et al. 2018; Li 2018) with the parameters -ax splice -uf -k10 -G2000.

### Estimating tail lengths using Nanopolish

We used Nanopolish v0.10.2 to estimate homopolymer tail lengths. First the fast5 and fastq files are indexed using Nanopolish index v0.11.1 with the default parameters. Then Nanopolish poly(A) v0.11.1 was used to estimate the length of the homopolymeric signal using the default parameters. We used the event detection module from nanopolish GitHub branch r10 for estimating signal variance for poly(I) and poly(A) homopolymers.

### Data deposition

Fast5 files acquired from nanopores used in this study have been uploaded to the European Nucleotide Archive https://www.ebi.ac.uk/ena/browser/home under the accession number PRJEB40734.

## Supporting information

Supplemental_Table_S2.xlxs

Supplemental_Table_S3.xlxs

## SUPPLEMENTAL MATERIAL

Supplemental Material is available for this article.

## ACKNOWLEDGEMENTS

Funding for this work was provided by NIH grants R01 HG010053 (Akeson) and R01 GM040478 (Ares). JQC was supported by a Jane Coffin Childs Foundation Postdoctoral Fellowship. Thanks to Craig Kaplan (University of Pittsburgh) for the tagged RNAPII yeast strain and advice on its use, and Olivia Rissland (University of Colorado, Denver) for advice on purifying recombinant Cid1. Thanks to Karla Neugebauer and her group for hosting a training session and sharing many details about their yeast chromatin isolation protocol.

