## Supplemental_Materials_Text_and_Figures.pdf for "Synthesis of modified nucleotide polymers by the poly(U) polymerase Cid1: Application to direct RNA sequencing on nanopores"

### Supplemental Figures

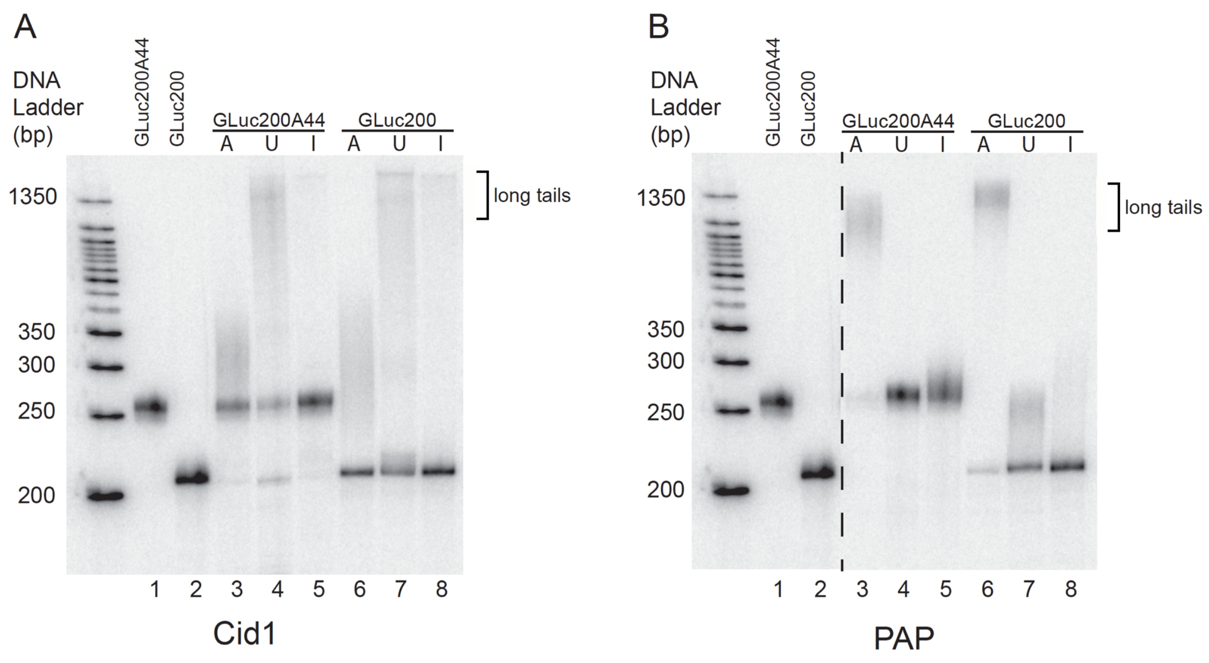


**Supplemental Figure S1**. Nucleotide adding activity of *S. pombe* Cid1 and *S. cerevisiae* Poly(A) polymerase (PAP) to the short model mRNA substrate Gluc200. **A**. Cid1 activity. **B.** PAP activity. RNA was incubated with enzyme and rNTPs as indicated and run on denaturing polyacrylamide gels, stained with SYBR Gold and imaged on a Typhoon scanner. Markers are DNA. Neither enzyme adds inosine very efficiently to either substrate. Apparently Cid1 is unable to add 50 inosine residues to the 44 nt polyA tail on the Gluc200(A)+ substrate although it does inefficiently form long I-tails on either substrate (panel A, lanes 5 and 8).

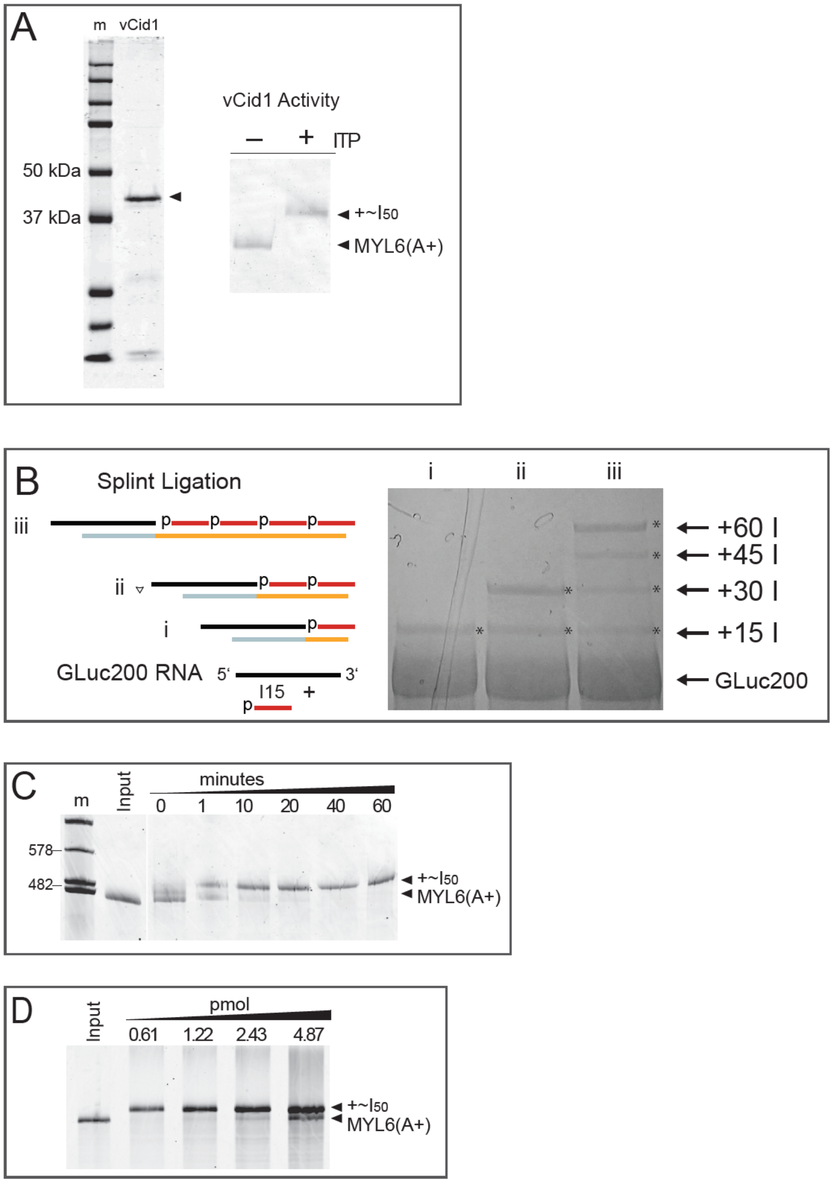


**Supplemental Figure S2.** Methods of addition of inosine to RNA. **A.** Homemade purified Cid1 expressed from *E. coli* adds 50 I residues to a model poly(A)+ mRNA. Left, a protein gel showing purified N-terminal truncated Cid1 (vCid1). Arrow indicates expected protein product (theoretical MW ~ 44.9 kDa). Right, inosine adding activity of purified vCid1 on MYL6(A)+ substrate showing major +50 nt product (+~$I_{50}$). **B.** Splint ligation was used to add defined lengths of $I_{15}$ to the end of a model mRNA. A different splint oligo (i, ii, or iii, blue and gold) was used to align the model RNA (black) and a phosphorylated inosine 15-mer (red) for ligation by T4 DNA ligase. The products were separated on a denaturing gel and gel purified. **C.** Time course of I-tailing on a model poly(A)+ mRNA. WIthin 20 min, addition of 50 I residues to all the substrate molecules is complete under the standard conditions (see Methods). **D.** Capacity of the standard reaction. Increasing amounts of substrate were I-tailed in a standard reaction (see Methods). Reaction capacity is about 2 pmoles under these conditions.

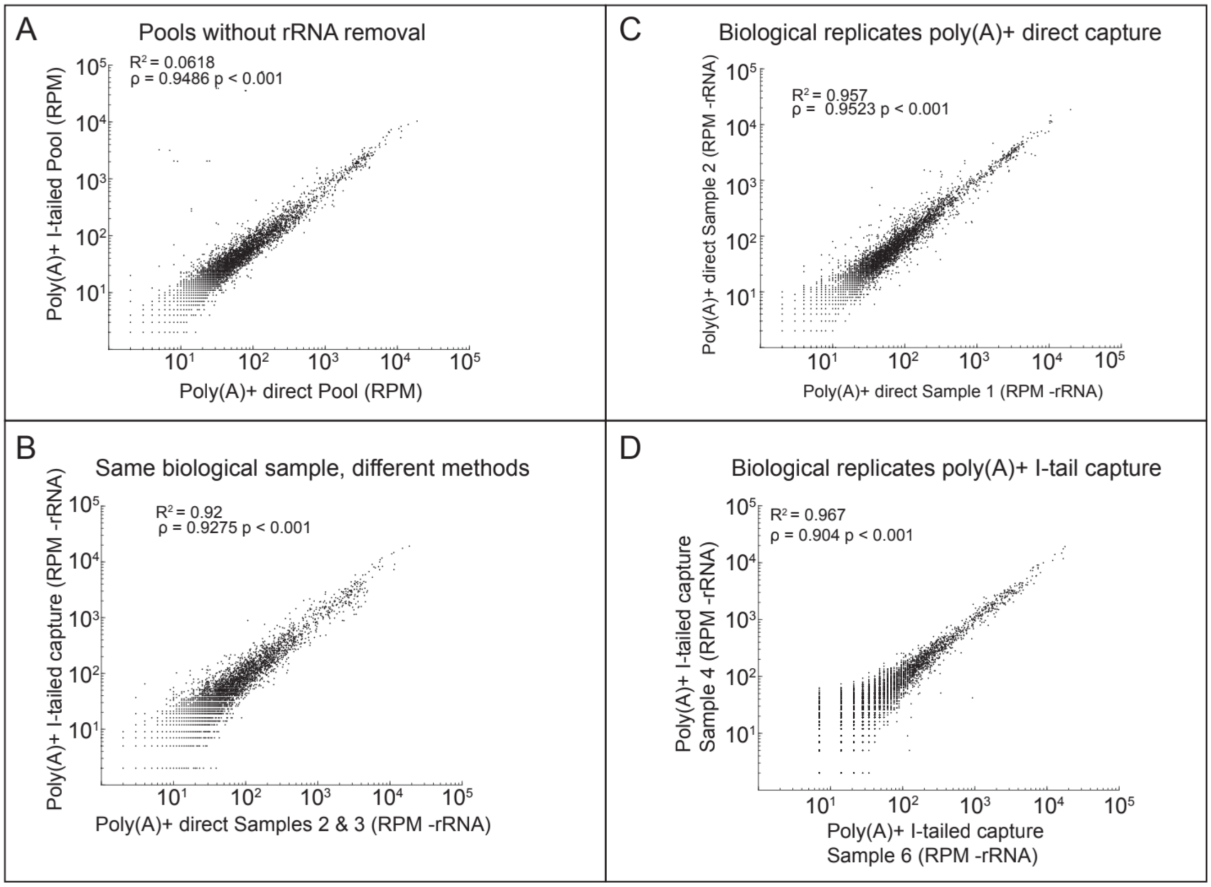


**Supplemental Figure S3.** Quantitative comparisons of libraries prepared by different methods. **A.** Poly(A)+ selected RNAs were either I-tailed and captured by the oligoC adapter and pooled together, or were captured directly by the standard oligoT and pooled together, then analyzed without removing rRNA reads. **B.** A single biological sample was analyzed by I-tailing or by the direct method. **C.** Two different biological replicates analyzed by the direct method. **D.** Two different biological replicates analyzed by the I-tailing method.

**Supplemental Table S1: Summary of sequencing runs**

| Library name | RNA treatment | Capture Method | # reads | # pass reads |
|---|---|---|---|---|
| Poly(A) Control #1 | Poly(A) #1 | ONT standard | 1,507,013 | 1,408,009 |
| Poly(A) Control #2 | Poly(A) #2 | ONT standard | 4,544,817 | 4,154,482 |
| Poly(A) Control #3 | Poly(A) #2 | ONT standard | 2,976,161 | 2,755,245 |
| Poly(A) Poly(I) #1 | Poly(A) #1 | Poly(I) | 496,472 | 435,203 |
| Poly(A) Poly(I) #2 | Poly(A) #1 | Poly(I) | 206,170 | 170,135 |
| Poly(A) Poly(I) #3 | Poly(A) #1 | Poly(I) | 347,347 | 302,453 |
| Poly(A) Poly(I) #4 | Poly(A) #3 | Poly(I) | 769,807 | 666,455 |
| Poly(A) Poly(I) #5 | Poly(A) #3 | Poly(I) | 34,427 | 26,291 |
| Poly(A) Poly(I) #6 | Poly(A) #3 | Poly(I) | 346,754 | 297,307 |
| Poly(A) Poly(I) #7 | Poly(A) #4 | Poly(I) | 367,679 | 319,259 |
| Total RNA | Total RNA (none) | Poly(I) | 128,919 | 57,276 |
| RiboMinus | RiboMinus | Poly(I) | 280,590 | 79,587 |
| RNAPII-associated | biotinylated-RNAPII streptavidin pull down | Poly(I) | 320,969 | 258,896 |

**Supplemental Table S2.** Supplemental_TableS2_Mapped_Reads.xlsx

**Supplemental Table S3.** Supplemental_Table3_Median_TailLengths.xlsx

### Supplemental Methods

#### Sequence of T7-MYL6 transcription template

This insert sequence (capital letters) is cloned in the SmaI site in the polylinker (lower case) of pUC13. The T7 promoter sequence is italicized. The recognition sequences of BsmI (GCATTC) and BbsI (GTCTTC) are underlined. The slash indicates the position of the 5’ end of the bottom strand of the cut DNA and is the last nucleotide transcribed by T7 RNA polymerase initiating at the T7 promoter, indicating the 3’ nucleotide of the MYL6(A)– transcript (BsmI cut template) and the MYL6(A)+ transcript (BbsI cut template).

gatcccc*TAATACGACTCACTATAGGGAG*ACAGTGGCCAAGAACAAGGACCAGGGCACCTATGAG
GATTATGTCGAAGGACTTCGGGTGTTTGACAAGGAAGGAAATGGCACCGTCATGGGTGCTGAA
ATCCGGCATGTTCTTGTCACACTGGGTGAGAAGATGACAGAGGAAGAAGTAGAGATGCTGGTG
GCAGGGCATGAGGACAGCAATGGTTGTATCAACTATGAAGCGTTTGTGAGGCATATCCTGTCGG
GGTGACGGGCCCATGGGGCGGAGCTCGTCCGCATGGTGCTGAATGGCTGAGGACCTTCCCA
GTCTCCCCAGAGTCCGTGCCTTTCCCTGTGTGAATTTTGTATCTAGCCTAAAGTTTCCCTAGGCT
TTCTTGTCTCAGCAACTTTCCCATCTTGTCTCTCTTGGATGATGTTTGCCGTC/A<u>GCATTC</u>ACCAA
ATAAACTTGCTCTCTGGAAAAAAAAAAAAAAAAAAAAAAAAAAAAAAAAAAAAAAAA/CT<u>GTCTTC</u>
gggcgagctcgaattc

**vCID1 expression in *E. coli*:**

Plasmid 10H-tev-vCID was made by standard techniques using synthetic DNA. It carries a beta-lactamase gene and a colE1 origin of DNA replication, and expression is driven by a T7 promoter. The predicted protein is 391 amino acids long with a theoretical molecular weight of 44.9 kilodaltons and the following sequence:

MGHHHHHHHHHHSSGAENLYFQSPNSHKEFTKFCYEVYNEIKISDKEFKEKRAALDTLRLCLKRISP
DAELVAFGSLESGLALKNSDMDLCVLMDSRVQSDTIALQFYEELIAEGFEGKFLQRARIPIIKLTSDTK
NGFGASFQCDIGFNNRLAIHNTLLLSSYTKLDARLKPMVLLVKHWAKRKQINSPYFGTLSSYGYVLM
VLYYLIHVIKPPVFPNLLLSPLKQEKIVDGFDVGFDDKLEDIPPSQNYSSLGSLLHGFFRFYAYKFEPR
EKVVTFRRPDGYLTKQEKGWTSATEHTGSADQIIKDRYILAIEDPFEISHNVGRTVSSSGLYRIRGEF
MAASRLLNSRSYPIPYDSLFEEAPIPPRRQKKTDEQSNKKLLNETDGDNSE*

To produce protein, *E. coli* BL21(DE3) pLysS cells are transformed with the plasmid and transformants are selected on LB agar plates with 100 µg/ml ampicillin at 37°C overnight. A single colony is used to inoculate 50mL of LB supplemented with 100 µg/ml ampicillin, shaken at 300rpm overnight at 37°C and diluted the next day into 1L with LB supplemented with 25 µg/ml ampicillin. After growth OD600 = 0.6 at 37°C and 300rpm, cells are induced to express vCID by adding IPTG to 1mM and shaking at 300rpm for 16-18 hours at 18°C. Cells are harvested by centrifugation at 5000rpm for 10 minutes, and cell pellets are resuspended in 10mL of 50mM Tris-HCl, 1mM EDTA pH 8.0 and centrifuged again at 4°C at 5000rpm, for 30 minutes. Supernatant is decanted, and washed pellets are stored at -80°C.

**vCID1 Purification**

Frozen cell pellets are resuspended in 14mL of a high salt-phosphate buffer (HSP buffer, 50mM sodium phosphate pH 8, 300mM NaCl, 100mM KCl, 1mM DTT, 10% glycerol) with 10 mM imidazole, with addition of lysozyme ot 1 mg/mL and PMSF to 0.5mM, and then incubated on ice for 5 minutes. Cells are lysed with glass beads by vortexing for 3 minutes in 15 second intervals (12 x 15 sec) with incubation on ice for 15 seconds between vortexing. Lysates are clarified by centrifugation at 14000 x g for 1 hour at 4°C. The resulting supernatant is incubated for 30 minutes with 2.5mL of cobalt-chelate resin (pre-equilibrated in 10 mM imidazole HSP buffer) with gentle shaking at 4°C, and then is poured into a column. The column is washed 2x with 12mL of 20mM imidazole HSP buffer, and then 1x with 5mL of 50mM imidazole HSP buffer. Eluates are collected using 100mM, 150mM, 200mM, 250mM, 400mM, and 500mM imidazole buffers at 5mL each in succession. Each eluted sample is concentrated in a protein concentrator tube (Millipore) until the total volume is <500uL. The samples are resuspended in 10mL storage buffer (10mM Tris-Cl pH 7.5, 100mM NaCl, 1mM DTT, 0.1mM EDTA, 50% glycerol v/v), and concentrated again until the sample volume was under 500uL.

**I15 Splint ligation for GLuc200I30 and GLuc200A44I30 library preparation**

To prepare GLuc200I30 and GLuc200A44I30 samples, 15pmol of GLuc200 RNA with 30pmol of the appropriate bottom splint adapter (C25TTTTTTTTTT (IDT) for GLuc200A44 or C25CCT AAG AGC AAG AAG AAG (IDT) for GLuc200), 1.4 nmols of a synthetic 5′p-15mer inosine homopolymer (Stanford PAN facility) in 10mM Tris-Cl pH 8.0, 1mM EDTA, and 50mM NaCl in 6ul reaction volume was heated to 55°C and slow cooled to 16°C in 25 minutes. Each reaction had 1ul 10X T4 ligation reaction buffer (NEB B0202S) and 2,000 units T4 DNA ligase (NEB M0202T) were added to each reaction and brought to 10ul volume with water then incubated at 16°C overnight. 2X RNA loading dye (2X NEB N0362) was added to each sample and denatured at 95°C for 5 minutes before loading into a 10% acrylamide gel and run for 3.5 hours at 28 watts. The gel excision was performed by post-staining with 1X SYBR Gold in TBE and visualized on a UV transilluminator while cutting with a razorblade. The samples were eluted from the gel slice in 850µl of 1x TAE buffer using a D-tubeTM Dialyzer Midi (Millipore Sigma 71507) for at least 90 minutes at 130 volts. The electro-eluted samples were precipitated with 85µl 0.3M NaOAc and 850µl isopropanol at -20°C overnight. The next day the samples were centrifuged at 4000 x g at 4°C for 30 minutes and the pellets were washed with 70% ethanol twice with subsequent centrifugations at 16000 x g for 15 minutes. The pellets were air dried for 15 minutes and resuspended in 10µl water with yields between 25-100ng. The libraries for each ligation product were prepared following ONT's Direct-RNA Nanopore sequencing library preparation with ~50ng of RNA using the polydC adapter. The optional reverse transcription step was skipped.
